# Flagellar energy costs across the Tree of Life

**DOI:** 10.1101/2022.01.31.478446

**Authors:** Paul E Schavemaker, Michael Lynch

**Affiliations:** Biodesign Center for Mechanisms of Evolution, Arizona State University, Tempe, AZ, 85287, USA

## Abstract

Flagellar-driven motility grants unicellular organisms the ability to gather more food and avoid predators, but the energetic costs of construction and operation of flagella are considerable. Paths of flagellar evolution depend on the deviations between fitness gains and energy costs. Using structural data available for all three major flagellar types (bacterial, archaeal, and eukaryotic), flagellar construction costs were determined for *Escherichia coli, Pyrococcus furiosus*, and *Chlamydomonas reinhardtii*. Estimates of cell volumes, flagella numbers, and flagellum lengths from the literature, yield flagellar costs for another ∼200 species. The benefits of flagellar investment were analysed in terms of swimming speed, nutrient collection, and growth rate; showing, among other things, that the cost-effectiveness of bacterial and eukaryotic flagella follows a common trend. However, a comparison of whole cell costs and flagellum costs across the Tree of Life reveals that only cells with larger cell volumes than the typical bacterium could evolve the more expensive eukaryotic flagellum. These findings provide insight into the unsolved evolutionary question of why the three domains of life each carry their own type of flagellum.

## Introduction

Life contains a dazzling diversity of molecular and cellular mechanisms. These are the consequence of, and are subject to, evolution. Though speculation abounds, there is yet no full integration of molecular and cellular biology with evolutionary theory (Lynch et al., 2014; Lynch & Trickovic, 2020). All features of the cell require energy for construction and operation, but they differ in their energy demand and contribution to fitness. Energy costs act as weights to evolutionary paths, if we want to know why only certain molecular and cellular features exist, we need an accounting of their baseline costs. Such an accounting has been under way for individual genes (Lynch & Marinov, 2015), membranes (Lynch & Marinov, 2017) and various other features (Lynch & Trickovic, 2020; Milo & Phillips, 2016b; Raven & Richardson, 1984). Many single-celled species swim by virtue of their flagella. However, why the three domains of life – bacteria, archaea, and eukaryotes – each evolved their own type of flagellum is still a major open evolutionary question.

The bacterial flagellum (Berg, 2003) consists of a helical protein filament, sometimes sheathed by a membrane (Geis et al., 1993), that is attached, via a hook region, to a basal body embedded in the cell envelope. The filament is assembled by progressive addition of monomers of the protein flagellin to the tip, reached by diffusion through a central channel in the filament. Its rotation is driven by a proton or sodium gradient (Ito & Takahashi, 2017). The archaeal flagellum, or archaellum, is also a rotating flagellum but differs from the bacterial one in two key respects. The filament, lacking a central channel, is assembled from the base, and its rotation is driven by ATP hydrolysis (Albers & Jarrell, 2018). The eukaryotic flagellum, or cilium, is completely different from the other two. Its diameter is an order of magnitude larger (Khan & Scholey, 2018); it bends rather than rotates; and the bending is caused by motor proteins arranged along the length of the flagellum. Most eukaryotic flagella have two single microtubules in the middle, surrounded by nine doublet microtubules. The microtubules are connected by protein complexes such as inner and outer dynein arms (King, 2016), nexin-DRC (Heuser et al., 2009), and radial spokes (Pigino et al., 2011). This whole protein complex, the axoneme, is always surrounded by a membrane.

Flagellar construction and operating costs have been estimated for the bacterium *Escherichia coli* (Macnab, 1996) and a dinoflagellate eukaryote (Raven & Richardson, 1984). For *E. coli*, the construction and operating costs are 2% and 0.1%, respectively, relative to total energy expenditure (Macnab, 1996). Others found an operating cost of 3.4% (assuming 5 flagella, a 100 Hz rotation rate, and a 1 h cell division time) (Ziegler & Takors, 2020). The relative construction cost for the dinoflagellate was estimated at 0.026%. The relative operating cost was 0.08-0.9%, compared to total cell maintenance cost (Lynch & Marinov, 2015; Raven & Richardson, 1984).

Here, following on previous work (Lynch & Trickovic, 2020), we provide a fuller accounting of construction costs for model bacterial, archaeal, and eukaryotic flagella, and extend these results to a range of bacterial and eukaryotic species using information on flagellum number, flagellum length, and cell volume retrieved from the literature. Drawing from additional estimates of the operating costs of flagella, these results are discussed in the context of swimming speed, nutrient uptake, growth rate, scaling laws, effective population size, and the origin of the eukaryotic flagellum.

## Results

### Energy costs of flagellum construction

Flagellar motility burdens the cell with costs of construction and operation. The former refers to the energy required for synthesis of the proteins and lipids that constitute the flagellum, which includes both direct and opportunity costs (Mahmoudabadi et al., 2019). The operating cost is the energy associated with rotating (bacteria and archaea) or beating the flagellum (eukaryotes).

Starting from the framework previously outlined (Lynch & Marinov, 2015), and ignoring protein turn-over (which is only a minor contribution to costs (Lynch & Marinov, 2015)), we estimate the construction cost per protein component (in units of ATP hydrolyses) as:

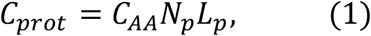

where *N*_*p*_ is the number of protein copies, *L*_*p*_ is the length of the protein in amino acids, and *C*_*AA*_ is the average energy cost of an amino acid (29 ATP; direct + opportunity costs (Lynch & Marinov, 2015)). When only the volume of a protein is known, we calculate the number of amino acids using the average volume per amino acid (1.33 × 10^−10^ μm^3^; Methods).

Eukaryotic flagella, and those of some bacteria, are enveloped by a membrane. Previously established approaches (Lynch & Marinov, 2017), with the inclusion of membrane proteins, leads to the membrane construction cost:

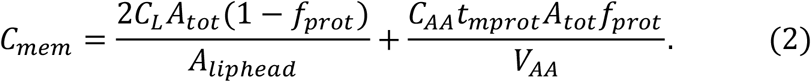

For the first term, *C*_*L*_ is the average energy cost of a single lipid molecule, *A*_*tot*_ is the total membrane surface area, which includes both lipids and membrane proteins, *A*_*liphead*_ is the cross-sectional area of a lipid head-group, *f*_*prot*_ is the fraction of membrane surface area occupied by protein (which is assumed to be 0.4) (Linden et al., 2012), and the factor 2 accounts for the bilayer of lipids. For the second term, *C*_*AA*_ is the energy cost per amino acid, *t*_*mprot*_ is the thickness of the protein areas of the membrane (8 nm), and *V*_*AA*_ is the average volume per amino acid.

Equations 1 and 2 yield estimates of *absolute* construction costs, in number of ATP’s. *Relative* construction costs are obtained by dividing the absolute flagellum costs by the construction cost of the entire cell (obtained from (Lynch & Marinov, 2015)). For a discussion of assumptions and omitted costs, see Methods. The flagellar construction cost data for the three model organisms are summarized in Table 1.

**Table 1:**
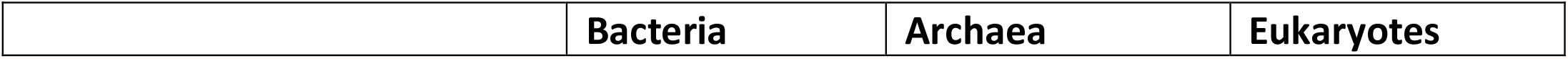
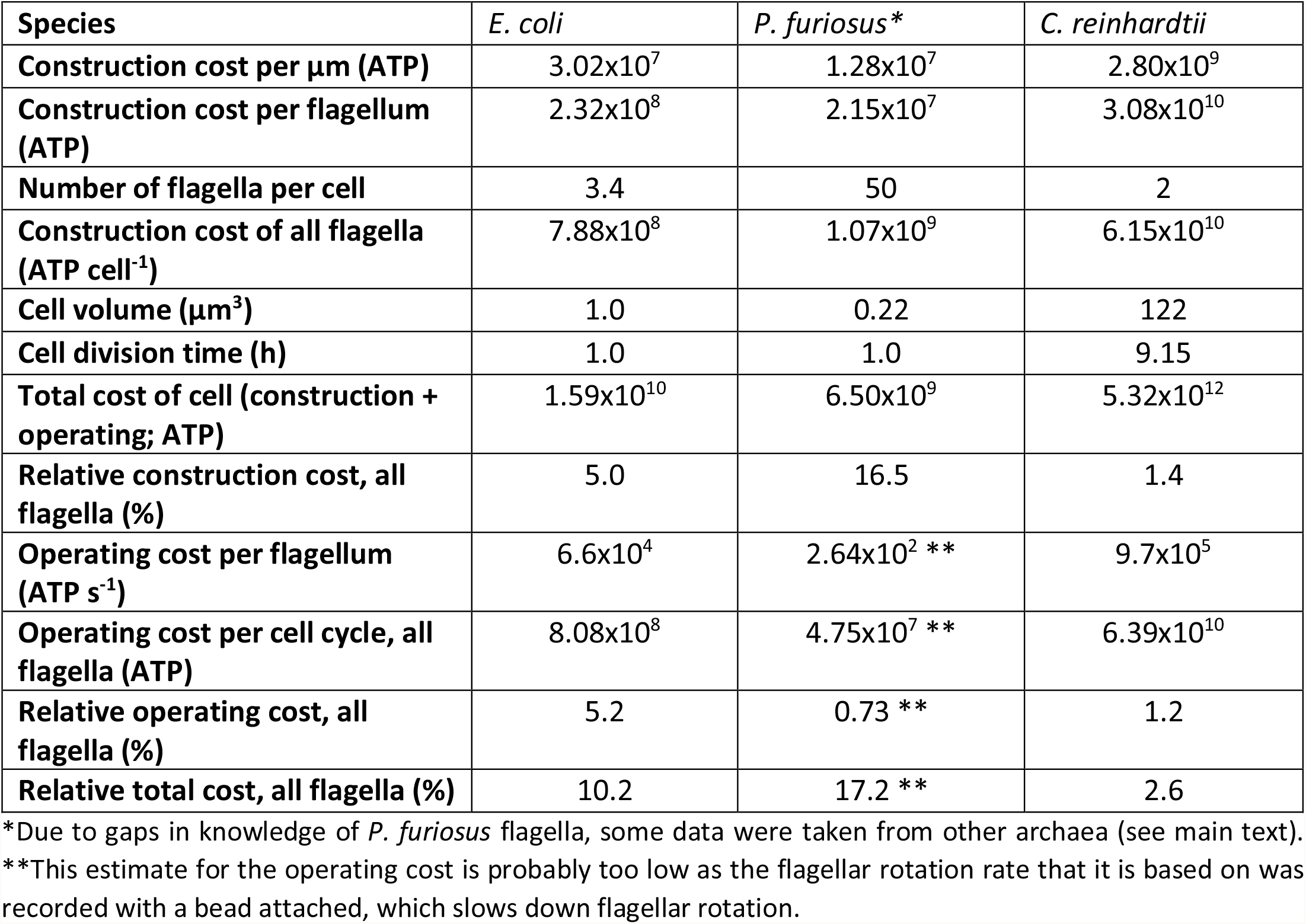
Energy costs of flagella in the three domains of life. Note that ATP denotes the number of ATP hydrolyses. A breakdown of the construction cost is available in Table 1-source data 1.

| Species | *E. coli* | *P. furiosus* * | *C. reinhardtii* |
| --- | --- | --- | --- |
| Construction cost per $\mu\text{m}$ (ATP) | $3.02 \times 10^7$ | $1.28 \times 10^7$ | $2.80 \times 10^9$ |
| Construction cost per flagellum (ATP) | $2.32 \times 10^8$ | $2.15 \times 10^7$ | $3.08 \times 10^{10}$ |
| Number of flagella per cell | 3.4 | 50 | 2 |
| Construction cost of all flagella (ATP cell<sup>-1</sup>) | $7.88 \times 10^8$ | $1.07 \times 10^9$ | $6.15 \times 10^{10}$ |
| Cell volume ($\mu\text{m}^3$) | 1.0 | 0.22 | 122 |
| Cell division time (h) | 1.0 | 1.0 | 9.15 |
| Total cost of cell (construction + operating; ATP) | $1.59 \times 10^{10}$ | $6.50 \times 10^9$ | $5.32 \times 10^{12}$ |
| Relative construction cost, all flagella (%) | 5.0 | 16.5 | 1.4 |
| Operating cost per flagellum (ATP s<sup>-1</sup>) | $6.6 \times 10^4$ | $2.64 \times 10^2$ ** | $9.7 \times 10^5$ |
| Operating cost per cell cycle, all flagella (ATP) | $8.08 \times 10^8$ | $4.75 \times 10^7$ ** | $6.39 \times 10^{10}$ |
| Relative operating cost, all flagella (%) | 5.2 | 0.73 ** | 1.2 |
| Relative total cost, all flagella (%) | 10.2 | 17.2 ** | 2.6 |

\*Due to gaps in knowledge of *P. furiosus* flagella, some data were taken from other archaea (see main text).
\*\*This estimate for the operating cost is probably too low as the flagellar rotation rate that it is based on was recorded with a bead attached, which slows down flagellar rotation.

### Bacteria – Cost of the *Escherichia coli* flagellum

For a detailed examination of the energy costs of the bacterial flagellum, we chose that of *Escherichia coli*. The protein composition of the flagellum, determined by a combination of structural and biochemical work, is summarized in (Berg, 2003). We supplemented this with copy numbers for export-apparatus proteins (Fukumura et al., 2017; Minamino, 2014, 2018) and verified the flagellin (FliC) copy number (Namba et al., 1989).

Using protein lengths and copy numbers, we calculated the energy costs for each protein per flagellum, and by summing these, obtained the cost for the whole flagellum. The total cost of a single 7.5 μm long (Turner et al., 2000) flagellum is 2.32 × 10^8^ ATP. A large fraction of this cost, 0.99, is in the filament (including the hook). With an average number of 3.4 flagella per *E. coli* cell (Harshey & Matsuyama, 1994; Turner et al., 2000), the total cost of flagella is 7.88 × 10^8^ ATP cell^-1^.

Because bacterial flagellum rotation is driven by the flow of protons from one side of the membrane to the other, an estimate of the operating cost requires information on the number of protons crossing the membrane, through the flagellum, per unit time. The energetic costs of this can be expressed in terms of the number of ATP hydrolyses by using the proton/ATP ratio in the ATP synthase, which is 3.33 (Jiang et al., 2001). The maximum number of stators, per flagellum, is at least 11 (Reid et al., 2006), each of which has two channels (Braun & Blair, 2001), and each channel passes 50 protons per revolution (Gabel & Berg, 2003; Samuel & Berg, 1996), leading to an estimated 1100 protons per revolution, which is similar to the value determined experimentally for *Streptococcus*, 1240 protons per revolution (Meister et al., 1987). The *E. coli* flagellum can spin at a maximal rate of 380 Hz (Chen & Berg, 2000; Gabel & Berg, 2003). The mean swimming speed for *Salmonella*, a close relative of *E. coli*, occurs at a rotation rate of 150 Hz (Magariyama et al., 2001). Taking a rotation rate of 200 Hz, a cell division time of 1 h, 3.4 flagella per cell, and assuming continuous operation we obtain a total operating cost of 1100×200×3.4×3600/3.33 = 8.08 × 10^8^ ATP cell^-1^.

For many evolutionary considerations, the cost of a structure/function relative to the cost of an entire cell is of interest. For *E. coli*, the construction (or growth) and operating (or maintenance) costs of the cell are estimated to be 1.57 × 10^10^ ATP and 2.13 × 10^8^ ATP h^-1^, respectively (Lynch & Marinov, 2015), and assuming a cell division time of 1 h implies a total cell energy cost of 1.59 × 10^10^ ATP. Thus, the costs of constructing and operating flagella relative to the whole-cell energy budget are 5.0 and 5.2%, respectively. The operating cost is close to an earlier estimate of 3.4% but differs from the other estimate of 0.1% (see introduction). How the 0.1% was obtained is not clear from the source.

Comparing the flagellum operating cost to just the cell *operating* cost leads to an apparent contradiction. The flagellar operating cost exceeds the cell operating cost by 3.8-fold. This mismatch could be due to the variability of *E. coli* cell volume (Taheri-Araghi et al., 2015) or the fact that the flagella are not continuously rotating. The cell operating cost may also have been determined under conditions in which cells do not swim. Temperature is probably not an issue as the whole-cell operating costs are normalized to 20°C (Lynch & Marinov, 2015), and the flagellum rotation speeds are determined at 23-24°C (Chen & Berg, 2000; Gabel & Berg, 2003).

### Archaea – Cost of the *Pyrococcus furiosus* flagellum

For the archaeal flagellum, we use the reasonably complete structural information on the flagellum of *Pyrococcus furiosus*, with some additional data from *Sulfolobus acidocaldarius* (Daum et al., 2017); see also (Albers & Jarrell, 2018). The major filament component in *P. furiosus* is FlaB0 (Nather-Schindler et al., 2014), with 1852 copies per μm of flagellum length (Daum et al., 2017) (extracted from PDBID: 5O4U). FlaB0 is chemically modified with oligosaccharides (35 sugar units per monomer) (Daum et al., 2017; Fujinami et al., 2014). We assume that each sugar unit costs as much energy as a single glucose in *E. coli*, which is 26 ATP (Mahmoudabadi et al., 2019). The length of the flagellar filament in *P. furiosus* is stated to be “a few 100 nm to several μm” (Daum et al., 2017). We assume this means 0.3-3 μm, or 1.65 μm on average.

Combining these data yields a construction cost of 2.15 × 10^7^ ATP per flagellum. As in *E. coli*, a large fraction of this cost is in the filament, 0.98. The number of flagella per *P. furiosus* cell is ∼50 (Daum et al., 2017), leading to a flagellar cost of 1.07 × 10^9^ ATP cell^-1^.

In addition to the flagella, *P. furiosus* has a polar cap that organizes all flagella into a tuft. The polar cap is rectangular and has a thickness of 3 nm and a linear dimension of 200-600 nm (Daum et al., 2017). Assuming a square of 400 × 400 nm implies a volume of 4.8 × 10^−4^ μm^3^. We also determined the total volume of a hexagonal array of protein complexes that is attached to the polar cap. Combining these volumes with the volume per amino acid we obtain a total polar cap construction cost of 2.43 × 10^8^ ATP cell^-1^.

The total cell cost is 6.50 × 10^9^ ATP cell^-1^, which is obtained by applying the *P. furiosus* cell volume (Daum et al., 2017) to the regression in (Lynch & Marinov, 2015) and assuming a 1 h cell division time. The relative cost for flagella plus polar cap is 20.2 % (16.5 % for flagella alone).

The rotation of the archaeal flagellum is driven directly by ATP hydrolysis, and 12 ATP are used for each rotation in *Halobacterium salinarum* (Iwata et al., 2019). The rotation rate is about 22 Hz (Iwata et al., 2019), but this may be an underestimate because in this measurement the flagellum was loaded with a 210 nm diameter bead, and in *E. coli* rotation rates decrease with an increased load (Gabel & Berg, 2003). We obtain an energy use of 264 ATP s^-1^ flagellum^-1^. Applying the *H. salinarum* numbers to *P. furiosus*, assuming a 1 h cell division time (Nather et al., 2006), yields an operating cost of 264×50×3600 = 4.75 × 10^7^ ATP cell^-1^.

*P. furiosus* has a peculiar metabolism and can grow at 100 °C (Kengen, 2017). We have not considered the impact of these differences on flagellar costs, but this should be investigated in the future.

### Eukaryotes – Cost of the *Chlamydomonas reinhardtii* flagellum

For the eukaryotic flagellum, we focus on *Chlamydomonas reinhardtii*, as its flagellum is very well characterised structurally. We include the axoneme, the membrane, and the intraflagellar transport system (IFT). The axoneme contains a central pair of microtubules surrounded by nine doublet microtubules. Each microtubule doublet consists of α- and β-tubulins and about 30 additional proteins (Ma et al., 2019). Bound to the outside of each doublet are the radial spokes (Pigino et al., 2011), the radial spoke stub (Barber et al., 2012), the IC/LC complex (Heuser et al., 2012), the CSC (Dymek & Smith, 2007; Pigino et al., 2011), the Nexin-DRC (Bower et al., 2013), the MIA complex (Yamamoto et al., 2013), the tether (Heuser et al., 2012), the outer dynein arms (King, 2016; King & Patel-King, 2015; Ma et al., 2019), and finally the inner dynein arms (King, 2013, 2016). The central pair of microtubules consist of α- and β-tubulins, and are bound by several other protein complexes (Carbajal-Gonzalez et al., 2013). To obtain the cost of the membrane, we assume a cylinder with a diameter of 0.25 μm (Khan & Scholey, 2018) and use Equation 2, with *C*_*L*_ = 406 ATP (Lynch & Marinov, 2017). Finally, we include the construction cost of 279 IFT complexes per flagellum (Vannuccini et al., 2016). Combining axoneme, membrane, and IFT, and assuming two flagella of 11 μm in length (Tuxhorn et al., 1998), we obtain a *Chlamydomonas* flagellar construction cost of 6.15 × 10^10^ ATP cell^-1^.

The operating cost of the *Chlamydomonas* flagellum determined experimentally on a naked axoneme (flagellum without the membrane) is 8.8 × 10^4^ ATP s^-1^ μm^-1^ (Chen et al., 2015). This is similar to estimates for a generic eukaryotic flagellum, 6.0 × 10^4^ ATP s^-1^ μm^-1^ (Raven & Richardson, 1984); *Paramecium caudatum*, 5.6 × 10^4^ ATP s^-1^ μm^-1^ (Katsu-Kimura et al., 2009) (Methods), assuming a total flagellum length of 1.70 × 10^5^ μm (Figure 1-source data 1); and sea urchin sperm, 6.2 × 10^4^ ATP s^-1^ μm^-1^ (Chen et al., 2015). The *Chlamydomonas* operating cost for a full 9.15 h cell cycle (Lynch & Marinov, 2015) is 6.39 × 10^10^ ATP.

The *Chlamydomonas* whole-cell construction and operating cost, at a 9.15 h cell division time, is 5.32 × 10^12^ ATP cell^-1^ (Lynch & Marinov, 2015), leading to the relative costs of constructing and operating both flagella of 1.4% and 1.2%, respectively. Comparing the flagellar operating cost (6.39×10^10^ ATP) to just the cell operating cost (2.4×10^11^ ATP per cell cycle) reveals that, unlike for *E. coli, Chlamydomonas* has enough energy in its operating budget to swim continuously.

### Flagellar construction costs across the tree of life

Flagellar construction costs for a wide range of bacterial and eukaryotic species were obtained by combining the just determined flagellar construction costs of *E. coli* and *Chlamydomonas* with flagellum lengths and numbers, and cell volumes for 196 species (27 Bacteria and 169 Eukaryotes) (Methods).

For bacteria, we assume that all flagella are built from flagellins of the same size as those in *E. coli* (FliC, 498 amino acids), the same holds for the other flagellar constituents. The absolute flagellar construction cost for bacteria spans 3.6 orders of magnitude, from 1.2 × 10^8^ to 4.4 × 10^11^ ATP (Figure 1A), with a median of 7.3 × 10^8^ ATP. The relative flagellar construction cost spans 2.3 orders of magnitude, from 0.43 to 96% (Figure 1B), with a median of 3.7%. The absolute cost of flagella increases with bacterial cell volume, but there is no discernible pattern relating relative costs and cell volume. A power law fit to the bacterial absolute cost vs cell volume data yields *c*_*abs*_ = 1.1 × 10^9^ *V*^0.81±0.14^ (exponent±SE) (with cell volume in μm^3^ and absolute cost in number of ATP hydrolyses).

**Figure 1.**
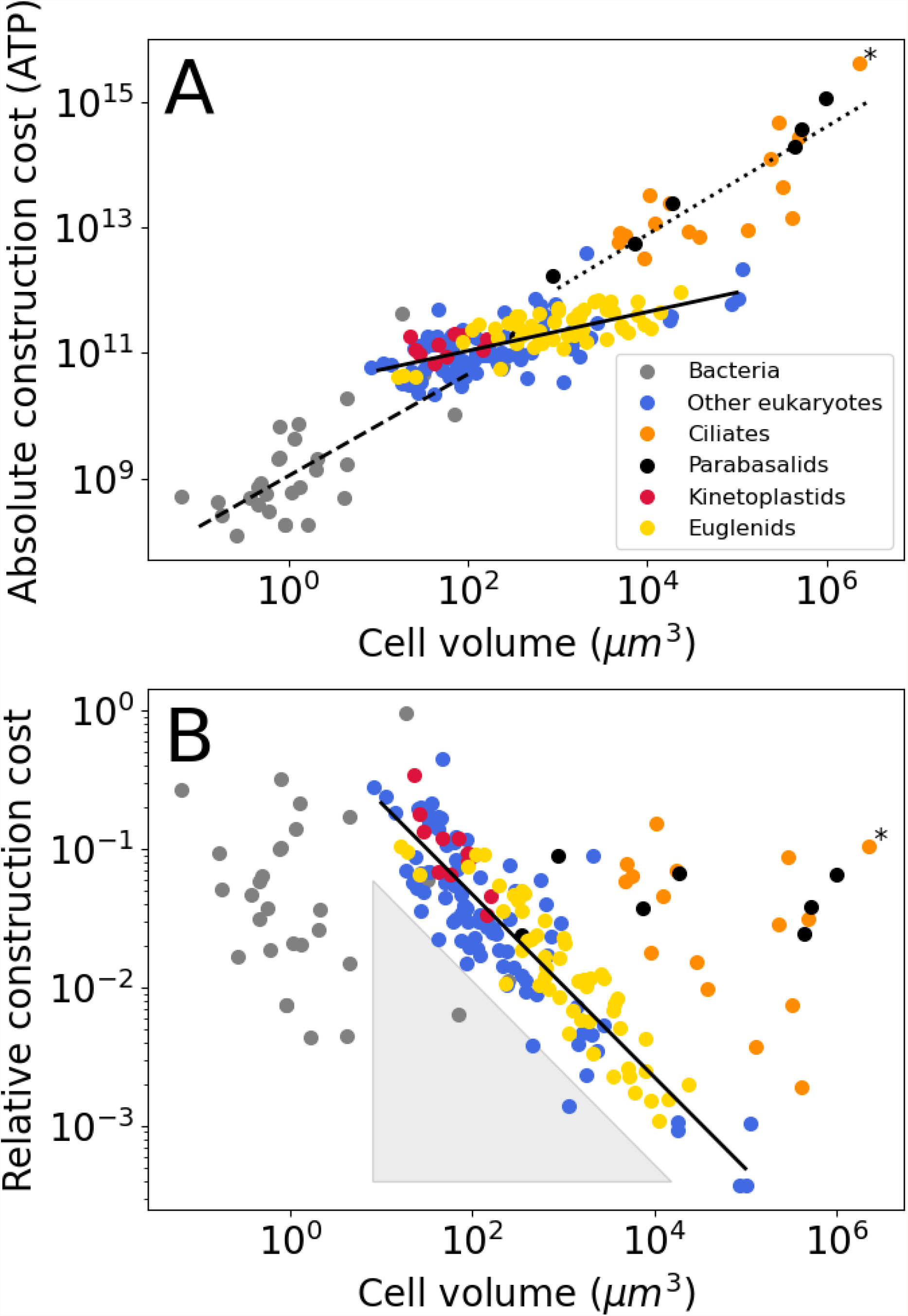
Construction costs of flagella in bacteria and eukaryotes as a function of cell volume. Each point denotes a species. The costs are the sum over all flagella present on a single cell. A) The absolute construction cost. The lines are power law fits to bacteria, eukaryotic flagellates (euglenids, kinetoplastids, and “other eukaryotes”), and ciliates and parabasalids. The equations are: *c*_*abs*_ = 1.1 × 10^9^ *V*^0.81±0.14^, *c*_*abs*_ = 2.6 × 10^10^ *V*^0.31±0.03^, and *c*_*abs*_ = 2.8 × 10^9^ *V*^0.86±0.10^, respectively (exponent ± SE). B) The construction costs of flagella relative to the construction costs of the entire cell. The line is a power law fit to the eukaryotic flagellates: *c*_*rel*_ = 0.98 *V*^−0.66±0.03^. The grey-shaded triangle is explained in the text. In both panels the asterisk marks *Opalina ranarum*, which resembles ciliates in its flagellar distribution but does not belong to the ciliate clade. Data in Figure 1-source data 1.

For most of the eukaryotes, we assume that flagella are of the same type as that of *Chlamydomonas*. For three phylogenetic groups, kinetoplastids, euglenids, and dinoflagellates, we also include the extra rod that they carry in their flagella besides the axoneme (Hyams, 1982; Maruyama, 1982; Portman & Gull, 2010; Saito et al., 2003), which increases the cost per unit length by twofold. We also assume, for all eukaryotes, that the flagellum maintains the same cross-sectional area throughout its length (though see Methods).

The absolute flagellar construction cost for eukaryotes spans 5.3 orders of magnitude, from 2.2 × 10^10^ to 4.2 × 10^15^ ATP (Figure 1A), with a median of 1.8 × 10^11^ ATP. The relative flagellar construction cost spans 3.1 orders of magnitude, from 0.037 to 44% (Figure 1B), with a median of 3.0%. Eukaryotic flagellates (euglenids, kinetoplastids, and other eukaryotes), carrying a small number of flagella, have lower absolute flagellum costs than the hyperflagellated ciliates and parabasalids. This is true even where cell volumes overlap. A power law fit to the eukaryotic flagellate absolute cost data has an exponent of 0.31±0.03, and a joint fit to the ciliates and parabasalids has an exponent of 0.86±0.10. In the plot of relative construction cost against cell volume, eukaryotic flagellates are again clearly separate from the ciliates and parabasalids. Here, the eukaryotic flagellates follow a power law scaling with an exponent of -0.66±0.03. An overview of eukaryotic flagellar and cellular traits is provided in Figure S1.

## Discussion

### Flagellar costs and benefits

To help understand why species evolve and maintain flagella, and why different domains of life rely on very different kinds of flagella, trait costs need to be compared to trait benefits. Here, benefits are analysed in three steps, from an increase in swimming speed to nutrient uptake to faster cell growth.

Swimming speeds were obtained from the literature for a subset of species from our cost dataset, revealing that swimming speed increases weakly with cell volume (power law exponent ±SE = 0.13±0.03) (Figure 2A), except perhaps for eukaryotic flagellates, which appear to have a volume-independent swimming speed, exponent = 0.02±0.09 (Figure 2A). There is, however, considerable noise in the data. Previous work, using a different set of organisms, suggested an exponent of 0.22±0.02 for bacteria and eukaryotes combined, and an exponent of 0.14±0.04 for eukaryotic flagellates (Lynch & Trickovic, 2020). Besides increasing with cell volume, the swimming speed also increases with absolute flagellar construction cost (exponent = 0.16±0.03; Figure 2B).

**Figure 2.**
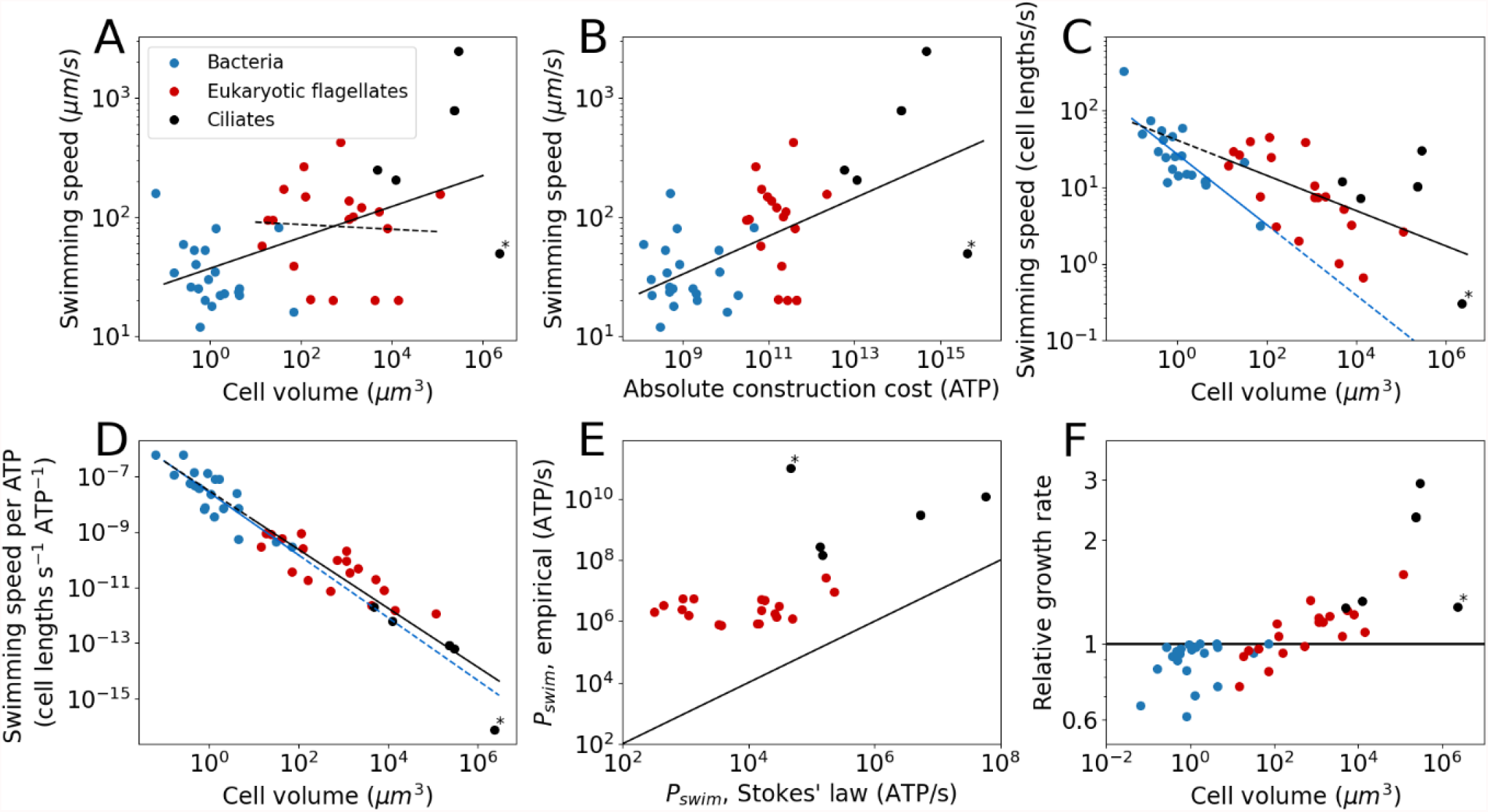
The cost and benefits of flagellar motility. Each point denotes a species. A) Swimming speed plotted against cell volume. Plotted are all species from the cost dataset for which the swimming speed is known. This includes bacteria, eukaryotic flagellates, and ciliates. The continuous line is the fit to all species: *v* = 37*V*^0.13±0.03^ (exponent ± SE). The dashed line is a fit to the eukaryotic flagellates: *v* = 95*V*^−0.02±0.09^. The legend holds for the entire Figure 2. B) Swimming speed plotted against the absolute flagellar construction cost. The line is the fit: *v* = 1.2*c*_*abs*_ ^0.16±0.03^. C) Swimming speed plotted against cell volume, but with swimming speed expressed in cell lengths per second (calculated from the cell volume by assuming a sphere). The solid blue line is the fit to the bacterial data: *v* = 27*V*^−0.46±0.08^. The solid black line is the fit to the eukaryotic data (both flagellates and ciliates): *v* = 41*V*^−0.23±0.07^. The dashed lines are extrapolations. D) Swimming speed (again in cell lengths/s) per ATP of construction cost plotted against cell volume. The solid blue line is the fit to the bacterial data: *v*_*ATP*_ = 2.7 × 10^−8^*V*^−1.13±0.16^. The solid black line is the fit to the eukaryotic data (both flagellates and ciliates): *v*_*ATP*_ = 3.1 × 10^−8^*V*^−1.06±0.11^. The dashed lines are extrapolations. E) Comparison of the swimming power, or operating cost, calculated from Stokes’ law with empirical values. This gives an indication of the efficiency of converting chemical energy into swimming power. The line indicates equality. F) The relative growth rate as a function of cell volume for cells in a medium with a homogenous distribution of small molecule nutrients, comparing cells with flagella to cells without flagella. In all panels the asterisk marks *Opalina ranarum*, which resembles ciliates in its flagellar distribution but does not belong to the ciliate clade. Data in Figure 2-source data 1.

Next, we compared bacterial and eukaryotic flagella in their ability to generate swimming speed, starting by plotting the swimming speed, normalized to cell lengths per second, against the cell volume (Figure 2C). From this plot, it may appear that the bacterial flagellum is more effective at small cell volumes, and the eukaryotic flagellum more effective at large cell volumes. However, this does not consider differences in how flagellar construction cost scales with cell volume in bacteria vs eukaryotes. Plotting the swimming speed per ATP invested against cell volume reveals that bacterial and eukaryotic flagella follow a common trend (Figure 2D), suggesting that there is no large difference in cost-effectiveness between the two groups once scaling with cell size has been taken into consideration. However, data for each flagellar type (bacterial or eukaryotic) only cover a limited, and mostly non-overlapping, volume range. Whether the common trend will be maintained over a broader range of cell volumes is unclear, but doubtful for the eukaryotic flagellum (see below).

To swim, a cell not only needs to construct a flagellum, but to operate it as well. We estimate operating costs in two separate ways. The first deploys Stokes’ law, which describes the power needed to move a sphere of radius, *r*, at a speed, *v*, through a medium with viscosity, *η*:

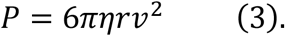

The second approach multiplies the ATP hydrolysis rate per μm of flagellum with the total flagellar length for each species (considering only eukaryotes). For the ATP hydrolysis rate, we used the average of the empirically determined values for Chlamydomonas, Paramecium, and sea urchin sperm (68700 ATP s^-1^ μm^-1^; see above). Comparing these two operating costs (Figure 2E) reveals that eukaryotes have a mean efficiency of 0.7% for the conversion of chemical energy (ATP) into swimming power, which is similar to earlier reports of swimming efficiencies for single-celled organisms (Chattopadhyay, 2006; Osterman, 2011; Purcell, 1997).

Faster swimming speeds can be used to increase nutrient uptake rate. The benefit of swimming for nutrient uptake, or the lack thereof, has been discussed by others, but without reference to the flagellar construction cost (Wan, 2021). However, the net fitness benefit also depends on what it costs to build and operate the flagella that propel a cell. To determine swimming gains, the amount of nutrients that is obtained by swimming needs to be compared to the amount of nutrients that is obtained by a stationary cell through diffusion. For a homogeneous nutrient distribution this is accomplished by calculating the Sherwood number, *Sh* (Guasto et al., 2012):

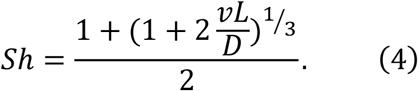

Here, *v* is the swimming speed (in μm/s), *L* is the cell diameter (in μm), and *D* = 10^3^ μm^2^/s is the small molecule diffusion coefficient (e.g. amino acids). A stationary cell has an *Sh* of 1, whereas an *Sh* of 1.1 means that swimming yields a 10% increase in nutrient uptake rate. For a flagellum to be selectively advantageous, the gain in nutrients must outstrip the cost of the flagellum. If the excess nutrients are used just for increasing the growth rate, then a relative growth rate is obtained as (Methods):

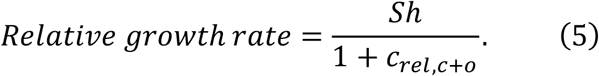

Here, *c*_*rel,c*+*o*_ is the relative flagellar cost, which includes both the construction and operating cost. It is assumed that the growth rate is nutrient limited and that for all species the flagellar operating cost is equal to the construction cost (Table 1). For species with a cell volume < 10^2^ μm^3^, flagella constitute a net loss in nutrients and decrease in growth rate (Figure 2F). This suggest that for these species, flagella cannot profitably be used for gathering nutrients in a homogenous medium. For species with a cell volume > 10^3^ μm^3^, flagella can provide a net gain in nutrients and therefore constitute an increase in growth rate.

### Scaling properties of flagellar energy costs and swimming speed

Here, we compare the empirical power-law relationships between flagellum construction cost, swimming speed, and cell volume (Figures 1B and 2A) with expected relationships derived from simple physical principles. If a simple model captures the general observed behaviour, considerations of morphological details and ecological backgrounds of each species and cells of different volumes become secondary issues. From Stokes’ law, already described above (Equation 3), we derived two power laws to compare with the empirical patterns in Figures 1B and 2A; detailed derivations can be found in the Methods.

The first power law concerns the relation between cell volume and relative flagellar construction cost observed for the eukaryotic flagellates in Figure 1B. Stokes’ law describes the power required for swimming, i.e. the flagellar *operating* cost. To make the connection to the flagellar *construction* cost in Figure 1B, it is assumed that flagellar operating cost and construction cost are linearly related (Methods). We also assume a spherical cell shape. A relative flagellar construction cost is obtained by dividing the absolute flagellar cost by the whole cell cost (which is obtained from the cell volume using an empirical relation (Lynch & Marinov, 2015)). Finally, based on the swimming speed data for eukaryotic flagellates (Figure 2A), we assume that swimming speed is independent of cell volume. These considerations yield (Methods):

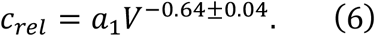

Here, *c*_*rel*_ is the relative flagellar construction cost, *a*_1_ is a constant, and *V* is the cell volume. The exponent in Equation 6, matches with the empirically determined exponent for eukaryotic flagellates (which excludes ciliates and parabasalids), -0.66±0.03 (Figure 1B). This suggests that the scaling of flagellar construction cost is determined by simple physical principles combined with the fact that swimming speeds do not vary with cell volume. The constant swimming speed is presumably the result of ecological factors. This result can be extended to any set of species in Figure 1B that are arranged along a diagonal with a slope of -0.64, so that, for example, all species along the bottom left edge of the ciliate distribution should also have the same swimming speed. Another corollary of this result is that there appears to be a (soft) lower limit to the swimming speed of eukaryotic flagellates, as there is a distinct lack of eukaryotic species in the grey shaded triangular area shown in Figure 1B.

The second power law concerns the relation between the cell volume and the swimming speed that is observed in Figure 2A (continuous line). Here, unlike the previous paragraph, we consider not just the eukaryotic flagellates, but the bacteria and ciliates as well. As above, we start from Stokes’ law and assume a spherical cell shape. We also assume that the operating cost is a fixed fraction of the total cell operating (or maintenance) cost. These considerations yield (Methods):

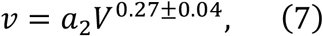

where *v* is the swimming speed, and *a*_2_ is a constant. The model exponent in Equation 7 is somewhat higher than the empirical result (Figure 2A; 0.13±0.03), as well as that from a previous analysis (Lynch & Trickovic, 2020), 0.22±0.02. This discrepancy may be due to the following. Cells with larger volumes tend to have many flagella. It may be that those flagella interfere with one another, reducing the efficiency of converting energy into swimming speed. This would lower the swimming speed for larger cells and therefore decrease the exponent (as compared to the model). Another possible cause may be the decreased relative energy investment in flagella in eukaryotic flagellates of larger cell volume (Figure 1B).

### Proteins from different parts of a flagellum experience different evolutionary forces

Flagella are composed of many different proteins which are present in a broad range of copy numbers. For example, a single *E. coli* flagellum contains one FliJ protein (part of exporter), ten FliD’s (filament cap), 100 FlgE’s (hook), and 10^4^ FliC’s (filament). The addition of a single amino acid to FliC has a larger impact on the cell energy budget than a single amino acid addition to FliJ, and thus requires a higher positive contribution to the cell fitness to be favoured by natural selection. Also, in the absence of any positive effect on fitness, a single amino acid addition to FliC is less likely to drift to fixation.

Here, we quantify the relative cost of adding a single (average) amino acid to selected flagellar proteins of all species for which we have data. For bacteria, the relative energy cost of adding an (average) amino acid to flagellar protein *i* in bacterial species *j* is:

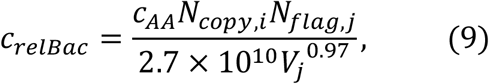

where *c*_*AA*_ is the average amino acid cost (29 ATP; (Lynch & Marinov, 2015)), *N*_*copy,i*_ is the protein copy number in a flagellum, *N*_*flag,j*_ is the number of flagella, and *V*_*j*_ is the cell volume. The denominator is the total cell construction cost (growth cost; (Lynch & Marinov, 2015)). For eukaryotes, which have a different flagellar architecture, we use a slightly different expression:

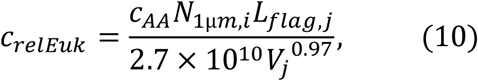

where *N*_1μ*m,i*_ is the protein copy number per μm of flagellum, and *L*_*flag,j*_ is the total flagellum length (the sum over the lengths of all flagella of a single cell). The resultant cost distributions (over all species) for selected flagellar proteins are shown in Figure 3.

**Figure 3.**
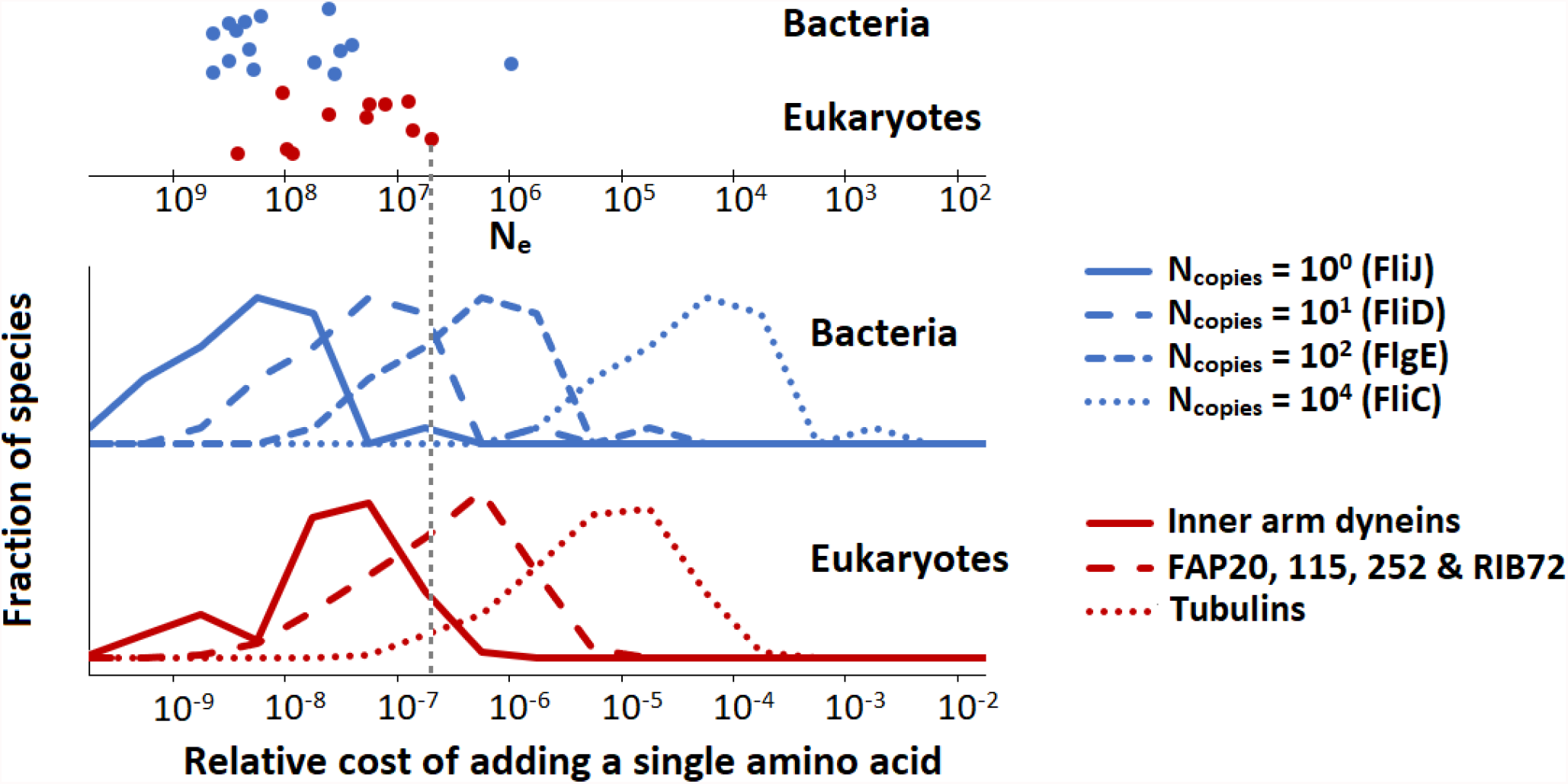
The population-genetic environment of different flagellar proteins for bacteria and eukaryotes. The different distributions are for different proteins, with varying copy numbers, within the same flagellum. The spread within each distribution reflects the variation of relative flagellar construction costs over all bacterial or eukaryotic species in our construction cost dataset. The points on the top show the effective population sizes, *N*_*e*_, of different species of bacteria and eukaryotes (Lynch & Trickovic, 2020) (the vertical spread of the datapoints is for visualisation). The bacterial flagellar protein names were taken from *E. coli*. Inner arm dyneins are present in 94 copies per μm of flagellum, for FAP20, etc., this number is 1125, and for tubulins it is 29125. The vertical dashed line is explained in the main text. Data in Figure 3-source data 1.

These data can be used to determine the minimal fitness benefit that an amino-acid addition needs to bestow upon the cell for the net fitness effect to be positive. We assume that for the small energy costs considered here, the relative energy cost is equal to the baseline loss in fitness so that the fitness before the amino acid addition is scaled to 1 and after the addition is 1 - *c*_*relBac*_ or *c*_*relEuk*_ (Ilker & Hinczewski, 2019; Lynch & Marinov, 2015; Lynch & Trickovic, 2020). If the amino acid is added to FliC, the *c*_*relBac*_ is roughly between 10^−6^ and 10^−3^ (depending on the species). The beneficial effect of the amino-acid addition needs to exceed this value by an amount exceeding the power of random genetic drift to be promoted by positive selection; and similarly, in the absence of any ecological fitness advantage, the insertion will be vulnerable to fixation by effectively neutral processes should the power of genetic drift exceed the cited values. For the low copy number FliJ, the comparable benchmarks (*c*_*relBac*_) are between 10^−10^ and 10^−7^, so the vulnerability to drift is increased by several orders of magnitude. The key point is that low copy-number flagellar proteins are expected to be more variable in length than those in high copy number, especially in species with small effective population sizes. This point is illustrated in Figure 3 with the vertical dashed line (indicating a species with an effective population size, *N*_*e*_, of 5×10^6^). Proteins with copy numbers that put them to the left of this line can accumulate amino acids neutrally, whereas the higher copy number proteins to the right of this line cannot. Similar considerations hold also for amino acid substitutions.

### Evolution of the eukaryotic flagellum

Archaea, bacteria, and eukaryotes each have their own type of flagellum. The reason for this independent evolution is an unsolved problem in evolutionary cell biology, made perhaps more important by the fact that the transition to a eukaryotic flagellum is part of the plethora of cellular modifications that led to eukaryotic cells. The three flagellar types are so different, in both assembly and thrust generating mechanisms, that it is hard to imagine a gradual transition between any of them. However, given that many prokaryotic and eukaryotic species do not possess a flagellum at all, the different flagellar types likely evolved *de novo*, in lineages initially devoid of flagella. A eukaryotic flagellum might also have evolved alongside a prokaryotic one, initially carrying out a separate function such as gliding or sensing. There are in fact bacterial species such as *Vibrio parahaemolyticus*, with two kinds of flagella, a thick one for swimming and multiple thin ones for gliding (both of the bacterial type) (McCarter, 2004).

For the many prokaryotic species lacking flagella, there may typically be enough food in the vicinity so that the costs of a flagellum outweigh its benefits, resulting in its loss. This may have been true also for the prokaryotic lineage that led to the eukaryotes. If so, the unique structure and mechanism of the eukaryotic flagellum may have owed more to contingency than to an advantage in swimming ability. However, eukaryotic cells also tend to be larger in volume than prokaryotic cells, and it is worth considering the consequences of this for the choice of flagellar type. For instance, it may not be possible for the thin prokaryotic flagellum to provide the thrust required to propel a large cell, as it would buckle under the load. This points to a possible advantage of possessing a thicker, and thereby stronger, eukaryotic flagellum. However, a few observations argue against this possibility. First, multiple bacterial flagella can be combined into a single large flagellum, as is the case for the magnetotactic bacterium MO-1 (Ruan et al., 2012). Second, a large bacterial cell could simply have many small flagella, as is the case for the giant bacterium *Epulopiscium* (Clements & Bullivant, 1991). Finally, there is a large single-celled eukaryotic species whose swimming is known to be powered by bacterial flagella, *Mixotricha paradoxa*, accomplished by recruiting bacterial species as motility symbionts (Wenzel et al., 2003). It remains to be determined whether the prokaryotic flagellum-powered motility of *Epulopiscium* and *Mixotricha paradoxa* can compete with the eukaryotic flagellum-powered motility of large eukaryotes. Combining flagellum construction costs with swimming speeds for a diversity of bacterial and eukaryotic species with a range of cell volumes (Figure 2D) suggests that there is no large advantage to having a eukaryotic flagellum in terms of the cost-effectiveness of generating swimming speed. However, a minor advantage cannot be ruled out.

In contrast, does the small cell volume of prokaryotic organisms prevent the acquisition of a eukaryotic flagellum? To answer this question, we first compare the absolute construction cost of entire bacterial cells to the construction cost of eukaryotic flagella (Figure 4A), revealing that the cheapest eukaryotic flagella are in fact more expensive than the entire cells of many bacterial species. Next, we examine the observed absolute flagellar investment in these bacterial species and ask what length of *eukaryotic* flagellum they could afford. These lengths are compared with the total flagellum length of eukaryotic species, revealing that most bacterial species cannot afford full length eukaryotic flagella (Figure 4B). Two observations suggest that the short and stumpy eukaryotic flagella these bacteria could afford would be highly ineffective. First, all eukaryotic flagella that have been observed, appear to be long and slender. Second, all small eukaryotes have a high relative investment in flagella (Figure 1B), suggesting that short (but cheaper) flagella are not up to the task. Reducing the cost of the eukaryotic flagellum by decreasing the number of microtubule doublets occurs in some species (Prensier et al., 1980; Schrevel & Besse, 1975), but its rarity suggests that it is associated with a considerable reduction in swimming ability.

**Figure 4.**
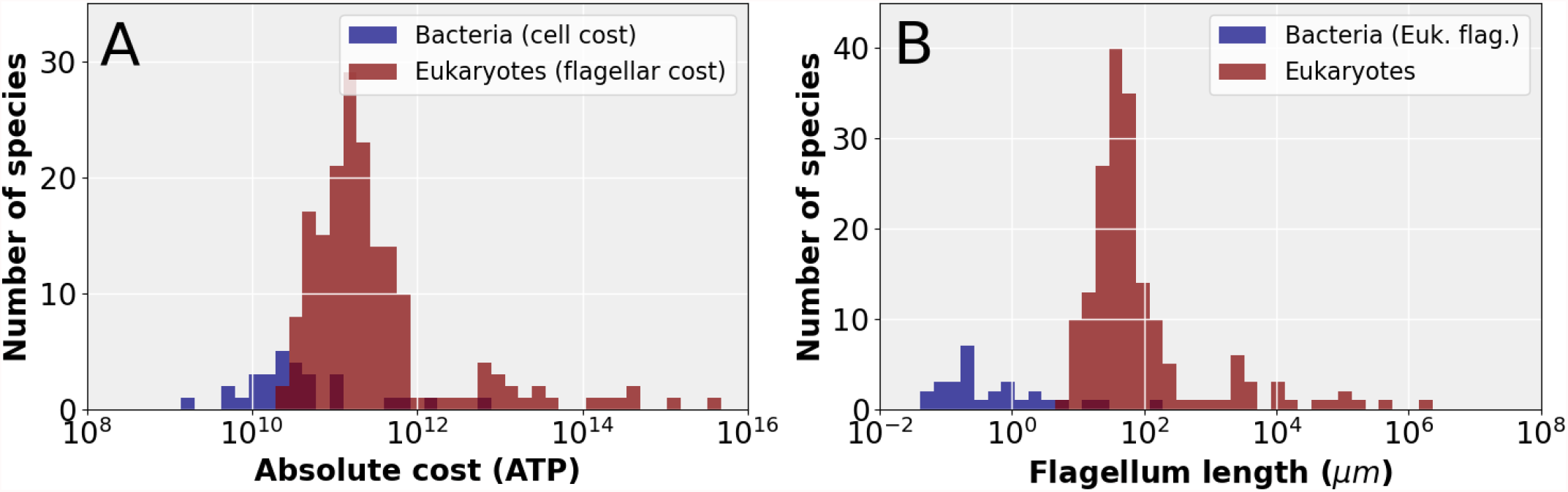
The cost of eukaryotic flagella compared to bacterial cell and flagellar budgets. A) Histograms comparing the total cellular cost (excluding maintenance) for bacterial species to the cost of constructing flagella in eukaryotic species. B) Histograms comparing the total *eukaryotic* flagellum length of bacteria and eukaryotes, where for the bacteria we have taken the absolute flagellar construction cost for each species and divided that by the per μm construction cost of the eukaryotic flagellum, to obtain the hypothetical length of the eukaryotic flagellum affordable to each bacterial species. Data in Figure 4-source data 1.

The possibility of an evolutionary origin of a reduced form of the eukaryotic flagellum in small cells cannot be ruled out. Consider the small alga *Micromonas pusilla* which has a cell volume of about 1 μm^3^ and a eukaryotic flagellum (Simon et al., 2017). Here, however, unlike most eukaryotic flagella, the outer microtubule doublets and motor proteins are only present in the proximal 10-20% of the flagellar length. The remainder of the flagellum consists of the inner pair microtubules surrounded by a membrane (Vaulot et al., 2008). Thus, the basic machinery that operates the eukaryotic flagellum, e.g. outer doublet microtubules, dynein motors, and radial spokes, may have evolved in a small prokaryote-sized cell; but the common eukaryotic flagellum, with this machinery present all along its length, could only evolve in larger cells.

## Conclusion

Using available structural data, we have determined the energy cost of building flagella in ∼200 unicellular species, including archaea, bacteria, and eukaryotes, recorded the swimming speeds for a subset of these, and reached the following conclusions. There appears to be a minimum swimming speed for eukaryotic flagellates, independent of cell volume. Flagella in small cells cannot be maintained solely for the collection of nutrients in homogenous media. Proteins from the same flagella but with lower copy numbers may experience a more permissive evolutionary environment, being able to accumulate amino acids neutrally and requiring lower beneficial effects for amino acid additions to be favoured by selection. There is no detectable difference in the cost-effectiveness of generating swimming speed between eukaryotic and prokaryotic flagella. Finally, the eukaryotic flagellum is only energetically affordable for cells that are larger than the typical prokaryote.

## Methods

### Data extraction

To calculate absolute and relative construction costs of flagella over a whole range of species, cell volumes and flagellar parameters were extracted from the literature. The cell volume was calculated from cell lengths (always excluding the flagellum), and widths, and occasionally also the cell depth. For bacteria a spherocylindrical cell shape and for eukaryotes a spheroidal cell shape was assumed. For some species, numbers for cell lengths, widths, and depths were not reported, so we extracted these from microscopy images, or, in some cases, from drawings.

The bacterial flagellum is helical but when a length is reported this could mean the arclength of the helix or the base to end distance. Wherever possible we used or calculated the arclength of the helix. When it was not clear what the reported length referred to, we assumed it was the arclength.

For ciliates and parabasalids, the number and length of flagella were in most cases not reported, so they were extracted from microscopy images and drawings. We made use of the regular spacing between flagella and the fact that they are arranged in rows and assumed that dikinetids have two flagella. Oral flagella were in most cases not included.

In ten cases there were more than one, and somewhat differing, datasets for the same species. These were included as separate points.

### Calculating the average volume per amino acid

The average molecular weight per amino acid residue is 110 g mol^-1^. The partial specific volume of proteins is 0.73 mL g^-1^ or 0.73 × 10^−6^ m^3^ g^-1^ (BNID 104272 and 110540)(Milo et al., 2010). Multiplying the amino acid molecular weight with the partial specific volume yields 80.3 × 10^−6^ m^3^ mol^-1^. Dividing this by Avogadro’s number gives 1.33 × 10^−28^ m^3^ residue^-1^ or 1.33 × 10^−10^ μm^3^ residue^-1^.

### Operating cost of Paramecium caudatum flagellum

The oxygen consumption rate of *P. caudatum* at different swimming speeds has been reported in the literature (Katsu-Kimura et al., 2009). This led to an estimate of the energy use for swimming at 1 mm s^-1^, which is 2.84 × 10^−6^ J h^-1^. The hydrolysis of ATP yields 5.0 × 10^4^ J mol^-1^ (Milo & Phillips, 2016a). From these numbers, and the fact that *P. caudatum* has 1.70 × 10^5^ μm worth of flagellum, operating cost per μm of flagellum length was determined: 5.60×10^4^ ATP s^-1^ μm^-1^.

### Flagellum assumptions

For the eukaryotic species that do not belong to the euglenids, kinetoplastids, or dinoflagellates, we treat the flagellum as if it were a *Chlamydomonas* flagellum, but with differences in length. This flagellum has a constant thickness throughout its length. For species that do belong to the euglenids, kinetoplastids, or dinoflagellates, the cost per unit length of flagellum was doubled to account for the presence of a rod. Here, we have also assumed constant thickness of the flagellum along its length. The constant thickness assumption appears to be borne out by the rod bearing *Trypanosoma brucei* (Portman & Gull, 2010), *Peranema trichophorum* (Saito et al., 2003), and *Ceratium tripos* (Maruyama, 1982) (*Ceratium tripos* is not in our dataset but *Ceratium fusus* is). We did find four species among the eukaryotic flagellates that have a flagellum that starts broad at the base and becomes thinner along its length, and that have a detailed enough description for us to make a cost calculation based on volume (assuming the same cost density as the *Chlamydomonas* flagellum). These species are *Anaeramoeba ignava, Anaeramoeba gargantua* (Taborsky et al., 2017), *Dinematomonas valida* (Larsen & Patterson, 1990), and *Psammosa pacifica* (Okamoto et al., 2012). In Figure S2 we plot the newly calculated relative cost as a function of the cell volume, as well as the old cost for comparison. This shows that there may be more overlap between the eukaryotic flagellates, and ciliates and parabasalids. Besides these four there appear to be more cells with somewhat thicker flagella at their base or throughout their length based on the drawings in (Larsen & Patterson, 1990; Patterson & Simpson, 1996), but not enough detail is given for us to perform any calculations.

Many eukaryotic flagella have hairs, either many along their length (mastigoneme) or one at the tip (acroneme), or vanes (Janouskovec et al., 2017; Tikhonenkov et al., 2014). As we do not know the detailed composition or exact size of these, we did not include their costs.

In calculating the relative costs of the flagella with respect to the total cell budget we have assumed that the flagella are expressed in every generation. If the flagella were expressed only once per ten generations, then the relative cost, in evolutionary terms, would be ten-fold lower.

Detailed structural information is only available for a limited number of eukaryotes. In principle there could be differences in the protein composition of the axoneme between different groups of organisms. A comparison of the *Chlamydomonas* flagellum with that of the ciliate *Tetrahymena* revealed only minor differences in axoneme structure (Pigino et al., 2012), which are inconsequential for our cost calculations. Some gametes are known to have 6+0 or 3+0 axoneme structure (Prensier et al., 1980; Schrevel & Besse, 1975), but these structures appear to be rare.

In our analysis of flagellar construction cost and swimming speeds we have assumed that all flagella were used for swimming. Flagella can also be used for gliding (Heintzelman, 2006) or capturing food. With more data on flagellar arrangements and descriptions of how flagella are in fact used, we will, in the future, be able to tease these different situations apart.

### Derivation of the relative growth rate equation

Here, we derive the relation of the relative growth rate to the Sherwood number, *Sh*, and the relative flagellar construction and operating cost, *c*_*rel,c*+*o*_ (Equation 5). We define the relative growth rate as:

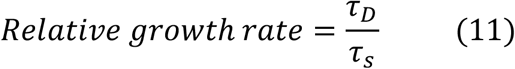

where *τ*_*D*_ is the cell division time in the absence of flagella and the supply of nutrients is by diffusion only, and *τ*_*s*_ is the cell division time in the presence of a flagellum and nutrient supply is increased by swimming. Assuming that cell division time is limited by nutrient supply, the cell division times can be calculated from the cost of the cell and the nutrient uptake rate.

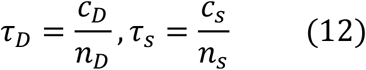

Here, *c*_*s*_ and *c*_*D*_ are the cell costs with and without the flagella, and *n*_*s*_ and *n*_*D*_ are the nutrient uptake rates with and without flagella. Equation 12 can be substituted into Equation 11 to obtain:

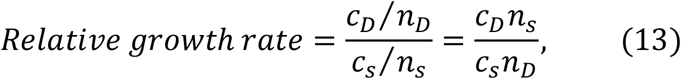

and since the Sherwood number is equal to the ratio of the nutrient uptake rates we find:

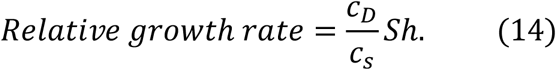

Next, the cell costs are normalized so *c*_*D*_ = 1 (just the cell) and *c*_*s*_ = 1 + *c*_*rel,c*+*o*_ (cell + flagella) yielding Equation 5.

### Derivation of scaling relations from Stokes’ law

Here we derive, under various assumptions, two power law relations from Stokes’ law: (1) a relation between cell volume and the relative flagellar construction cost in eukaryotic flagellates, and (2) a relation between cell volume and swimming speed. Stokes’ law describes the power, *P*, that is needed to propel a sphere of radius, *r*, through a liquid of viscosity, *η*, at a speed, *v*:

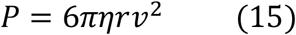

Since we are dealing with spheres, we can also express Equation 15 in terms of volume, *V*, as:

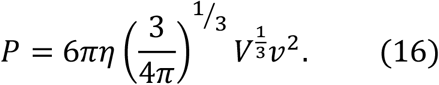

The first power law we derive is for the relation between cell volume and relative flagellar construction cost that is observed for the eukaryotic flagellates in Figure 1B. We do so for the special case in which the swimming speed doesn’t change with cell volume. This allows us to simplify Equation 16 to:

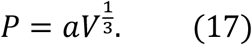

Here, *a* is used to indicate that there is a proportionality constant. It has no particular value and can’t be compared between, or within equations. We assume that the absolute construction cost, *c*_*abs*_, of a flagellum is linearly related to its power required for swimming (see below), so we have:

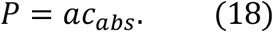

The absolute flagellar construction cost is related to the relative flagellar construction cost by (Lynch & Marinov, 2015):

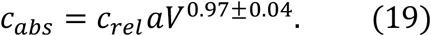

Now we combine Equations 17-19 to obtain the relation between cell volume and relative flagellar construction cost (Equation 6).

The reasoning behind flagellar operating cost being linearly proportional to flagellar construction cost is as follows. In the case of the eukaryotic flagellum the flagellum length is linearly proportional to the number of motors. So, a doubling of flagellar length (which doubles the construction cost), either by having two flagella or by doubling the length of the existing flagellum, also doubles the energy consumption. In the case of bacteria, the same argument can be made if doubling of construction cost means having two flagella, which would also double energy consumption. The effect of doubling the length of a bacterial flagellum is not so obvious. However, doubling the length of the bacterial flagellum presumably doubles the amount of water that is displaced per rotation, requiring a doubling of the operating cost. Interactions between flagella or between parts of the same flagellum, either directly or hydrodynamically, probably affect the operating cost. So, the linearity between operating and construction cost should be viewed as an approximation. There is also limited empirical support for the linear proportionality of flagellar operating and construction cost. The operating cost per μm flagellum in *Chlamydomonas* and *Paramecium* is similar (Results), despite them having a wildly different number of flagella.

The second power law we derive is for the relation between the cell volume and the swimming speed that is observed in Figure 2A. We start with Equation 16, and substitute the power, *P*, by the flagellar operating cost, *c*_*oper*_:

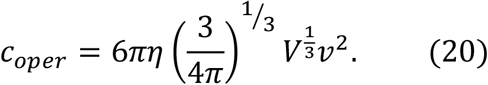

The flagellar operating cost is assumed to be a fixed fraction of the total cell operating (or maintenance) cost (see below), so that we can substitute the operating cost by a constant times the total cell operating cost, *c*_*oper,cell*_:

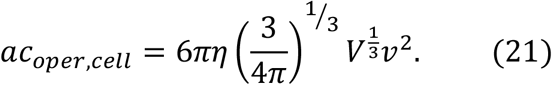

The total cell operating cost is related to the cell volume (Lynch & Marinov, 2015), so we have:

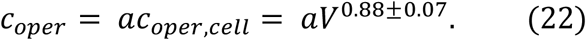

Combining Equations 21 and 22 yields:

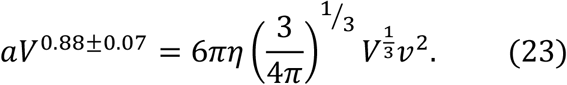

Simplifying this yields the relation between cell volume and swimming speed (Equation 7).

We want to compare Equation 7 to the data in Figure 2A. However, Equation 7 only holds when the flagellar operating cost, *c*_*oper*_, is a fixed fraction of the total cell operating (or maintenance) cost. This is true if both the flagellar operating cost and the total cell operating cost scale with cell volume in the same way. Which is what we will demonstrate here. However, we don’t have the flagellar operating cost data over a range of cell volumes. So, instead, we start by assuming that the flagellar *operating* cost is linearly proportional to the flagellar *construction* cost (justified above). Next, we determine the power law relation between cell volume and absolute flagellar construction cost for the subset of species for which we also have swimming speed data (all species in Figure 2A, but only a subset of the species in Figure 1A). This power law has an exponent of 0.86±0.05. We compare this to the empirically determined exponent in the power law relation between cell volume and total cell operating cost (Lynch & Marinov, 2015), 0.88±0.07. Because the exponents are the same, we can conclude that the flagellar construction cost, and thereby the flagellar operating cost, is indeed a fixed fraction of the total cell operating cost. Thus, it is safe to compare Equation 7 to the data in Figure 2A.

## Acknowledgements

We thank Sergio A. Muñoz-Gómez and Bogoljub Trickovic for critically reading and commenting on the manuscript. This work was supported by the US Department of Army, MURI award W911NF-14-1-0411; the National Institutes of Health, R35-GM122566-01; the National Science Foundation, DEB-1927159; and the Moore–Simons Project on the Origin of the Eukaryotic Cell, Simons Foundation 735927, https://doi.org/10.46714/735927.

---

**Figure S1.**
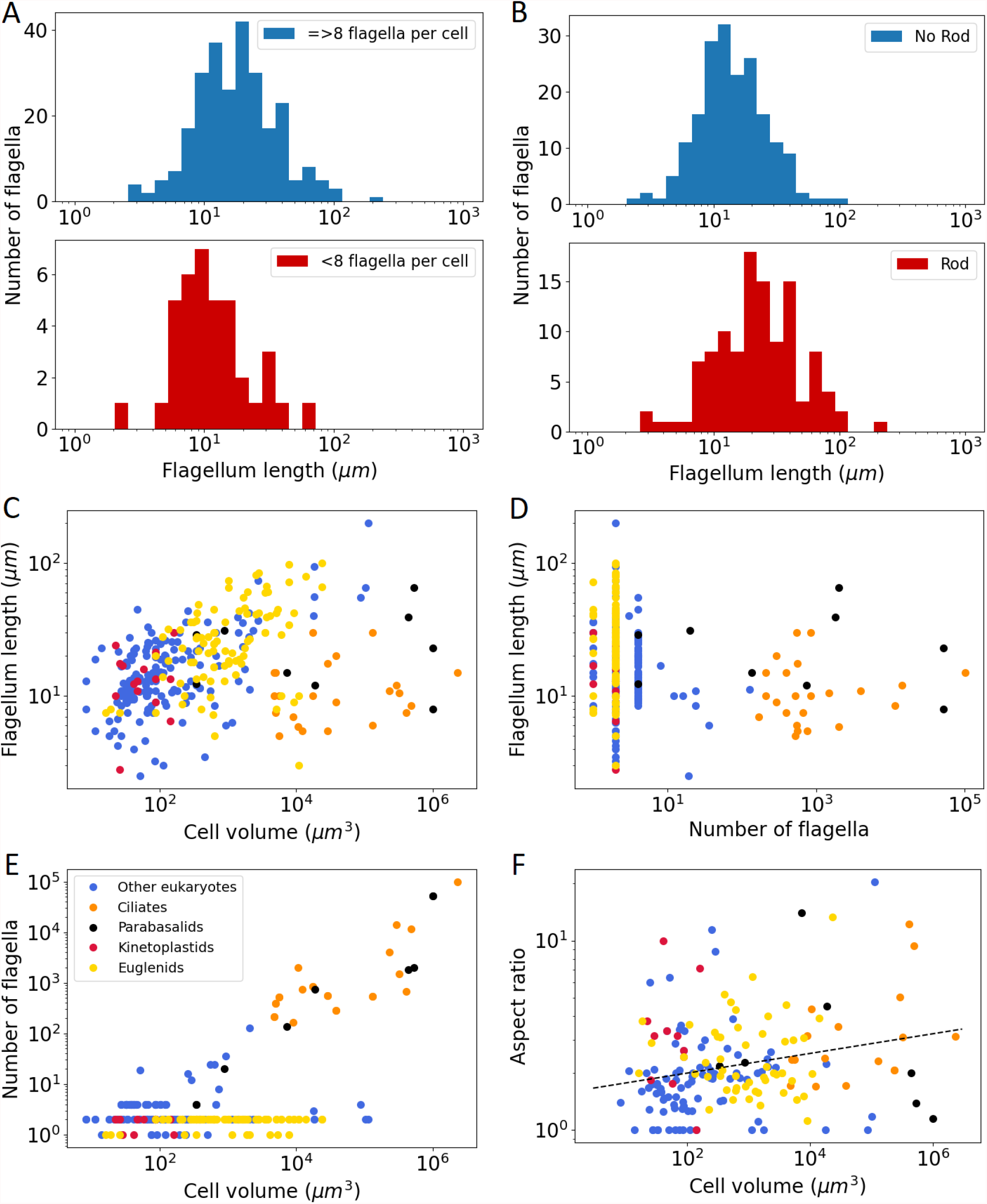
Overview of flagellar and cellular properties of eukaryotic species. A) Distribution of flagellum lengths for cells with less than eight flagella and cells with eight or more flagella. Only unique flagellar lengths for each species are included (also for Figures S1B-E), otherwise ciliate and parabasalid data would overwhelm the plot. B) Distribution of flagellum length for cells with or without a rod in their flagellum. C) Flagellum length plotted against cell volume. The legend in figure S1E holds for figures S1C-F. D) Flagellum length plotted against number of flagella. E) Number of flagella plotted against cell volume. F) Cell aspect ratio plotted against cell volume. The aspect ratio is calculated by dividing the cell length by the cell width (or the geometric mean of the width and depth). The dashed line is a power law fit to all species: 1.57 *V*^0.052±0.016^. Data in Figure S1-source data 1 and Figure S1-source data 2.

**Figure S2.**
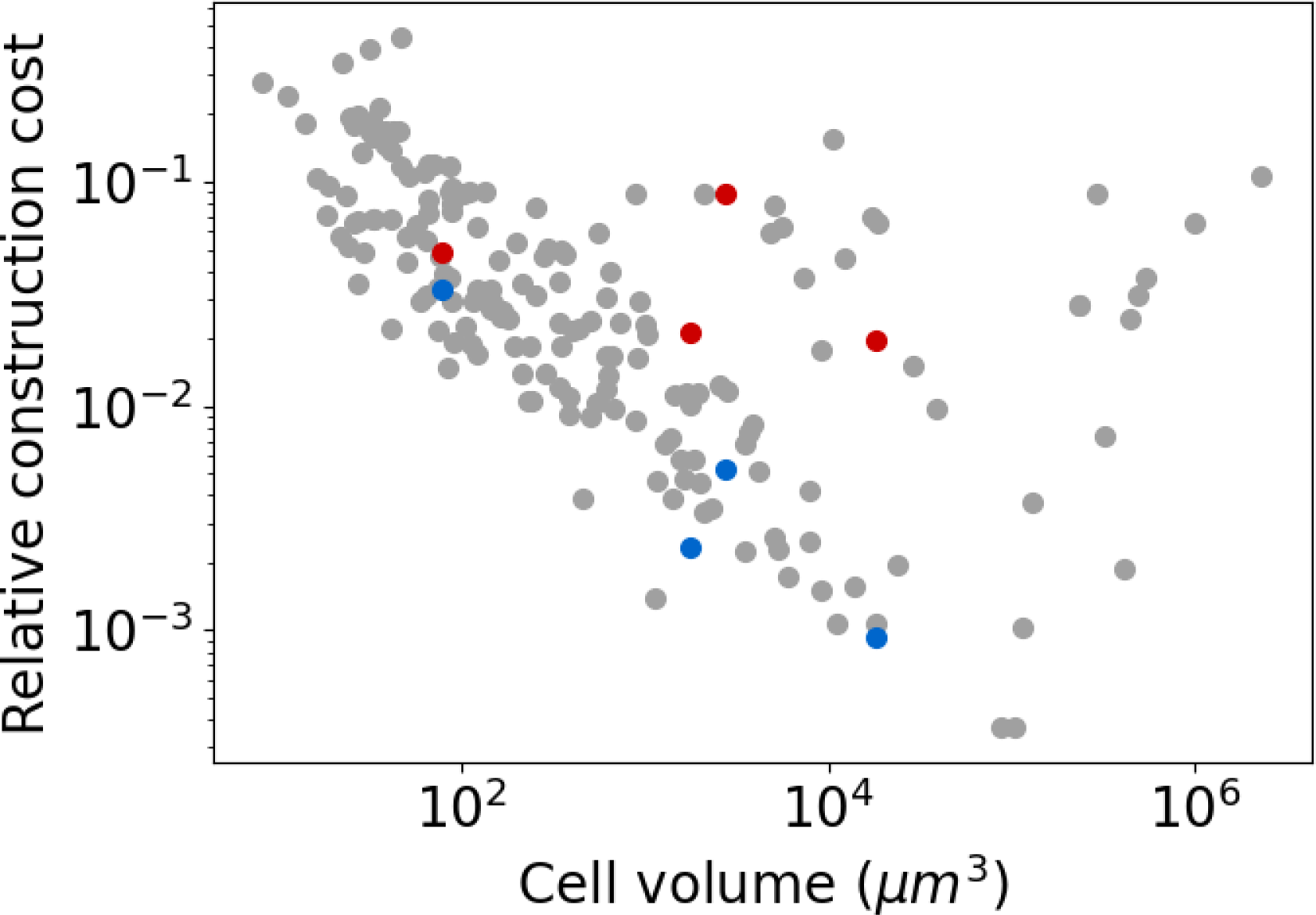
Relative flagellar construction costs of special cases plotted against cell volume. The blue points show *Anaeramoeba ignava, Anaeramoeba gargantua, Dinematomonas valida*, and *Psammosa pacifica*, before correcting for their real flagellum shape. The red points show the same species after correction. The grey points represent other eukaryotic species, shown for comparison. Data in Figure S2-source data 1.

